# Linking flexibility of brain networks to cognitive development in preschool children

**DOI:** 10.1101/2020.03.24.005074

**Authors:** Lily Chamakura, Syed Naser Daimi, Katsumi Watanabe, Joydeep Bhattacharya, Goutam Saha

**Affiliations:** Department of Electronics and Electrical Communication Engineering Indian Institute of Technology, Kharagpur, India; Department of Psychology Goldsmiths, University of London, London, United Kingdom; School of Fundamental Sciences and Engineering, Waseda University, Tokyo, Japan

**Author notes:** Corresponding Author Email addresses (Lily Chamakura), (Syed Naser Daimi), (Joydeep Bhattacharya), (Goutam Saha).

**Keywords:** brain network, dynamic functional connectivity, early childhood, individual differences, connectome predictive modeling

## Abstract

Recent studies of functional connectivity networks (FCNs) suggest that the reconfiguration of brain network across time, both at rest and during task, is linked with cognition in human adults. In this study, we tested this prediction, i.e. cognitive ability is associated with a flexible brain network in preschool children of 3-4 years - a critical age, representing a ‘blossoming period’ for brain development. We recorded magnetoencephalogram (MEG) data from 88 preschoolers, and assessed their cognitive ability by a battery of cognitive tests. We estimated FCNs obtained from the source reconstructed MEG recordings, and characterized the temporal variability at each node using a novel path-based measure of temporal variability; the latter captures reconfiguration of the node’s interactions to the rest of the network across time. Using connectome predictive modeling, we demonstrated that the temporal variability of fronto-temporal nodes in the dynamic FCN can reliably predict out-of-scanner performance of short-term memory and attention distractability in novel participants. Further, we observed that the network-level temporal variability increased with age, while individual nodes exhibited an inverse relationship between temporal variability and node centrality. These results demonstrate that functional brain networks, and especially their reconfiguration ability, are important to cognition at an early but a critical stage of human brain development.

## 1. Introduction

The preschool years, often considered as the “blossoming period” Brown and Jernigan (2012), is a critical phase in human development because it is associated with the most dynamic and global changes in the brain’s structure and functions Casey et al. (2005), yet our understanding of the functional brain network patterns in preschool children is limited. This period is characterized by the onset of executive functions Diamond (2006), memory Simcock and Hayne (2003) and most importantly, development of key skills such as language Bannard et al. (2009); Skipp et al. (2002) and reading Lonigan et al. (2000). The structure and function of the developing brain are closely related to behavior as well as cognitive and life outcomes Siugzdaite et al. (2020). For example, the maturation of the prefrontal cortex drives the rapid development of cognitive flexibility, a core component of executive functions, in the preschool years Buttelmann and Karbach (2017). Of note, cognitive flexibility is related to academic achievement (e.g mathematics or reading skills) Yeniad et al. (2013) and even temperament Quiñones-Camacho et al. (2019) in preschool children. Besides, neuroimaging studies of preschool children reveal associations between distinct brain activation patterns and individual differences in behavior Quiñones-Camacho et al. (2019) and cognitive abilities Cantlon and Li (2013). Predicting developmental and cognitive profiles from functional brain networks in preschoolers would be critically relevant as these profiles are often related to children’s learning ability Siugzdaite et al. (2020), and this is a principal aim of our current study.

The relevance of large scale whole-brain functional connectivity is widely established Bressler and Menon (2010). For example, functional brain network patterns in adults can predict individual differences in sustained attention Rosenberg et al. (2016), creativity Beaty et al. (2018) and personality traits Tompson et al. (2018); Markett et al. (2018) and further could distinguish between health and disease Bassett and Bullmore (2009). This network view assumes stationarity of the functional interactions while disregarding the temporal dynamics; pairwise functional connectivity (FC) is usually computed over the entire data duration (be it rest or task-related), leading to the estimation of a static network. Yet brain responses are transient in nature, so static connectivity estimate smooths out the variations across time and the resulting static network is widely different from the network at any of the time instants Allen et al. (2012). Importantly, numerous studies in adults have shown evidence for the dynamic nature of FC characterized by the variations of connection strengths, the sign of interactions or changes in node membership among modules Hutchison et al. (2013) during task conditions as well as rest Calhoun et al. (2014). Further such transient nature of brain network is reported to increase over development Chai et al. (2017), and directly related with learning Bassett et al. (2011), task performance Shine et al. (2016); Keerativittayayut et al. (2018), executive functions Medaglia et al. (2018), and in general, with healthy cognitive functioning Thomas Yeo et al. (2011) including general intelligence Barbey (2018) and creativity Li et al. (2017); Sun et al. (2018) in adults.

Compared to studies in adults, research on the dynamic reconfiguration of the typically developing brain is just beginning. For example, functional MRI (fMRI) analyses in a cohort of healthy children and young adult participants (3 - 23 years) revealed that the variability in the functional topography could predict individual differences in brain maturity Cui et al. (2020). In healthy children and adolescents (6-17 years), dynamic FC analyses of resting-state fMRI revealed diverse functional brain states as compared to static FC patterns Marusak et al. (2017); the participants also exhibited an age-related increase in variability suggesting greater neural complexity with maturation Marusak et al. (2017). Recent studies on children with autism spectrum disorder reported decreased variability of the default mode network (age group 3-7 years) He et al. (2018) and longer dwell times in disconnected global states (age group 6-10 years) Rashid et al. (2018) as compared to typically developing children.

Based on these studies, we hypothesized that dynamic FC could predict individual differences in cognitive abilities in preschool children. We investigated the dynamic FC networks (FCNs) of 88 preschoolers (36 - 59 months old) using Magnetoencephalogram (MEG) recordings acquired while they watched cartoon videos. The children’s cognitive abilities were assessed by Kaufman Assessment Battery for Children (K-ABC) tests Kaufman and Kaufman (1983), a widely used test battery for the assessment of intelligence and achievement in young children Sotel-Dynega and Dixon (2014); Benson et al. (2019). In particular, we had the following predictions (a) the dynamic FCN in children would be supported by mesoscale variations of the neural regions that exhibit diverse functional roles across time, (b) the extent of variation associated with each region would be modulated by its centrality in the network, (c) the temporal variability of the dynamic FCN would increase with age in the preschool years, and (d) the deviations of functional states in the dynamic FCN from the static FCN can predict performance on children’s cognitive abilities.

## 2. Materials and Methods

### 2.1. Participants

In total 88 preschool children (43 girls) with an age range between 35-59 months were recruited for this study. All children were native Japanese children without previous or existing psychological or developmental problems as verified by parents’ reports. All children were tested over two separate days; on the first day, they performed cognitive tests (K-ABC) and were familiarized with the MEG environment, and the actual MEG recording was performed on the second day. On both testing days, experimenters played with children along with the parents in order to ensure that the children felt comfortable and at ease in the laboratory. All experimental procedures were fully explained to the parents before they agreed, in written informed consent, to the participation of their child. The experimental protocol was approved by the Ethics Committee of Kanazawa University Hospital and the experiment was conducted in accordance with the World Declaration of Helsinki.

### 2.2. MEG recording

Magnetoencephalogram (MEG) was recorded with a 151-channel SQUID (superconducting quantum interference device) whole head coaxial gradiometer MEG system specialized for children (PQ1151R; Yokogawa/KIT, Kanazawa, Japan) in a magnetically shielded room. The custom child-sized MEG system allows easy placement and effective positioning of the sensors so that head movement is appropriately constrained (Johnson et al, 2010). The head position was determined by measuring the magnetic fields after passing currents through coils placed at three locations on the head surface as fiducial points with respect to well-defined locations (bilateral mastoids and nasion). During MEG recording, children were lying on the bed in a supine position while watching a cartoon of their own choice that was selected before the recording. The narration sound was delivered binaurally through a tube leading to speakers placed in front of the children. An experimenter remained close by to ensure the comfort of the children and further to prevent movement during the MEG recording. MEG data were sampled at a rate of 1 kHz and filtered with a 200 Hz low-pass filter.

### 2.3. K-ABC scores

The children were administered the Kauffman Assessment Battery for Children (K-ABC) test Kaufman and Kaufman (1983), which is a standardized test, based on the theoretical foundation of Luria’s Luria (1966) to measure cognitive development in children between 2.5 to 12.5 years of age. The K-ABC includes assessments of both intelligence and achievement, and is grounded in the theoretical foundation of test was conducted separate from the MEG recording. The participant scores were recorded for four scales - Sequential processing scale, Simultaneous processing scale, Mental processing scale and Achievement scale. In sequential/successive processing, the stimuli are processed in a sequence, and the stimuli have an inherent temporal order. The sub-tests for the sequential processing scale include word order (touching series of objects in same order as named by the examiner), number recall (repeating a sequence of numbers as said by the examiner) and hand movements (copying a sequence of taps on the table as performed by the examiner with palm/hand). On the other hand, in simultaneous processing, all pieces of information are available at once, which are integrated concurrently to arrive at a task solution. The sub-tests for simultaneous processing scale include triangles (assembling of colored rubber triangles to match an abstract image), gestalt closure (meaningful interpretation of partially completed pictures) and face recognition (recognizing the face shown by the examiner in a different (group) photograph). The sequential and simultaneous scales characterize short-term memory and visual-spatial abilities respectively. The mental processing scale is a composite of sequential and simultaneous processing scales where the latter has a greater influence. The mental processing composite score is considered the global estimate of a child’s level of intellectual functioning. The achievement scales measure achievement and focus on applied skills and facts that were learned from the school or home environment.

### 2.4. Preprocessing

MEG preprocessing was done using the FieldTrip toolbox Oostenveld et al. (2010) in MATLAB. For each participant, the MEG data in each channel was band-pass filtered between 0.5 Hz and 45 Hz using a 4th order Butterworth filter. The data was two-pass filtered to avoid phase distortions. Independent Component Analysis (ICA) Hyvärinen and Oja (2000) was used to removing eye blink and heartbeat artifacts. The bad channels were identified by visual inspection of the data and then interpolated using cubic splines from the neighboring channels determined through Delaunay triangulation. The 3-minute long recording was again visually inspected and bad segments were discarded. We discarded MEG data from a child if the duration of artefact free segment was less than 80 s, and this led to a loss of 14 participants, therefore leaving 74 MEG dataset (39 girls) for our analysis. For each participant, the preprocessed MEG was partitioned into *K* segments of *L* = 80 sec duration (here, segments were generated with 50% overlap between successive segments). Here, *K* depends on the total artifact-free data available and hence, varies across participants between 1 to 3. In the rest of the paper, we refer to the multiple segments per participant as sessions.

### 2.5. Source reconstruction

The source reconstruction was performed using FieldTrip toolbox Oostenveld et al. (2010). For source analysis, forward models were constructed using age-specific average MRI templates of 3-4 years old children from Neurodevelopmental MRI database Sanchez et al. (2012). The forward model was computed separately for 3 years and 4 years age groups using age-specific average MRI templates from the database. For beamformer solution, the bandwise covariance matrix was calculated by considering the pre-processed data of all subjects. The dipoles were assumed to be located at the voxels within the head boundary (only grey matter was considered) on a 3D grid with 5 mm spacing. This resulted in 9023 and 9710 voxels for 3 and 4 years age-groups respectively, where sources were to be localized. In each of the six frequency bands, the source activity was reconstructed first band-pass filtering the sensor data in that band and determining the beamformer filter coefficients through Linearly Constrained Minimum Variance (LCMV) approach Van Veen et al. (1997). The cortical reconstructions were parcellated into regions of interest (ROI) using according to the LONI Probabilistic Brain Atlas (LPBA40) atlas Shattuck et al. (2008). This resulted in a total of 56 cortical regions of interest with 27 identical regions in each hemisphere (see Supplementary Table S1) for list of regions along with their abbreviations). Subsequently, the first eigenvariate of reconstructed source activity was computed in each ROI to obtain the representative time series. The first eigenvariate was computed by projecting the reconstructed source activity in each ROI onto the first eigenvector after singular vector decomposition (SVD). Hence, in each band, we obtained a total of 56 time series corresponding to all cortical ROI in the source space.

### 2.6. Functional connectivity estimates

Functional connectivity (FC) estimates quantify the interdependence between brain activity at spatially distinct locations. In this study, we used *coherence* to estimate the pairwise functional connectivity between the source reconstructed time series. As the source space does not exhibit volume conduction effects, coherence yields reliable estimates of connectivity. The static and dynamic connectivity was computed in six frequency bands — Delta (0 - 4 Hz), Theta (4 - 8 Hz), Alpha (8 - 12 Hz), Beta-1 (12 - 20 Hz), Beta-2 (20 - 30 Hz) and Gamma (30 - 45 Hz) bands. Next, we briefly describe the computation of static and dynamic FC matrices.

#### 2.6.1. Computation of static connectivity

The static connectivity was estimated for each session using *Coherence* between all pairs of ROI. To estimate the static connectivity in each band, the MEG data was filtered according to the frequency band of interest using a band-pass filter (4^*th*^ order Butterworth filter). Let *x*_B_(*t*) and *y*_B_(*t*) denote the filtered data of the ROI pair under consideration whose Hilbert transform is denoted as 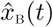 and *ŷ*_B_ (*t*) respectively. The corresponding analytic signals were then obtained as:

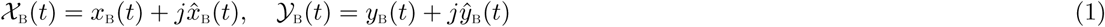

Then, the pairwise connectivity *C*_XY_ for the frequency band under consideration Cohen (2014) was obtained as:

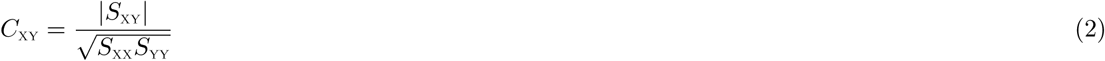

where 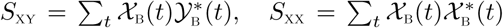 and 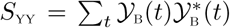 are scalars and denote the cross- and auto-power spectral densities in the considered frequency band *B*. Coherence takes values between 0 and 1; *C*_XY_ = 0 denotes that the time-series are linearly independent, while *C*_XY_ = 1 denotes perfect linear dependence. Since there are 56 ROI after parcellation, we obtain an undirected network with weighted adjacency matrix of size 56 × 56.

#### 2.6.2. Computation of dynamic connectivity

The time-varying connectivity was estimated by partitioning each session of the MEG data into overlapping windows and computing the coherence (equation 2) in successive time-windows. This sliding window approach requires the choice of the resolution (determined by window size) and overlap parameters. The topology of the network depends (among other factors) upon the time-scale at which it is defined Horwitz (2003). The window resolution must be fine enough to detect short duration events. However, choosing very small window sizes is not feasible because it can lead to insufficient number of data points for estimating the power spectra and in turn connectivity. For this reason, we constructed multiple temporal networks for each session with different window sizes of *M* = 2.5, 5, 7.5, 10, 12.5 and 15 sec, with 50% overlap between successive windows.

### 2.7. Computation of temporal variability

Here, we define the temporal variability of the nodes with respect to their dissimilarity to their role in the static network. For each node, the proposed measure captures the deviation of the dynamic FC states, which may reflect task-relevant network topologies, from the static FC. For node *i* of the dynamic FCN, it is defined as:

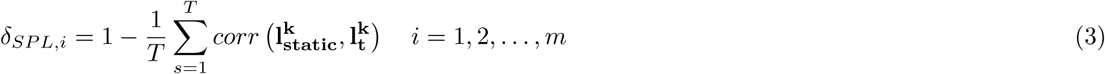

where corr(.) denotes the Spearman rank correlation of the vectors; 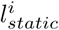 is the (*m* − 1)-dimensional vector comprising the shortest path lengths of node *i* to the rest of the nodes in the static network; 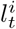 is the vector of shortest path lengths during the time-window indexed by *t* in the temporal network. The measure *δ*_*SPL,i*_ captures the extent to which the nodes differ from their average behavior as captured by the static functional network. Higher the value of *δ*_*SPL,i*_, higher the variability of node *i*. To obtain a feature value per ROI per participant, we averaged the values across multiple sessions and window sizes for each participant.

The proposed measure in equation 3 is based on the measure of nodal temporal variability introduced by Zhang et al. (2016). It is computed as below for node *i* in the temporal network:

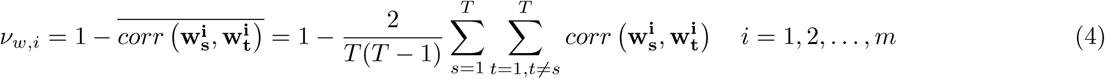

Here, **w**_**t**_^*i*^ = [*w*_*ij*_]_*j*≠*i*_ denotes (*m* − 1) dimensional vector corresponding to the node *i*, comprising the connection weights from *i* to all other nodes at time *t*. Further, *corr*(.) denotes Spearman rank correlation between vectors. The measure in equation 4 suffers from one significant drawback. Since the functional architecture of the node at any instant is represented by its connection weights to other nodes (**w**_**t**_^*i*^), it fails to capture the relevance/role of the node in context of the whole network. We illustrate this with an example in Figure 1A.

**Figure 1:**
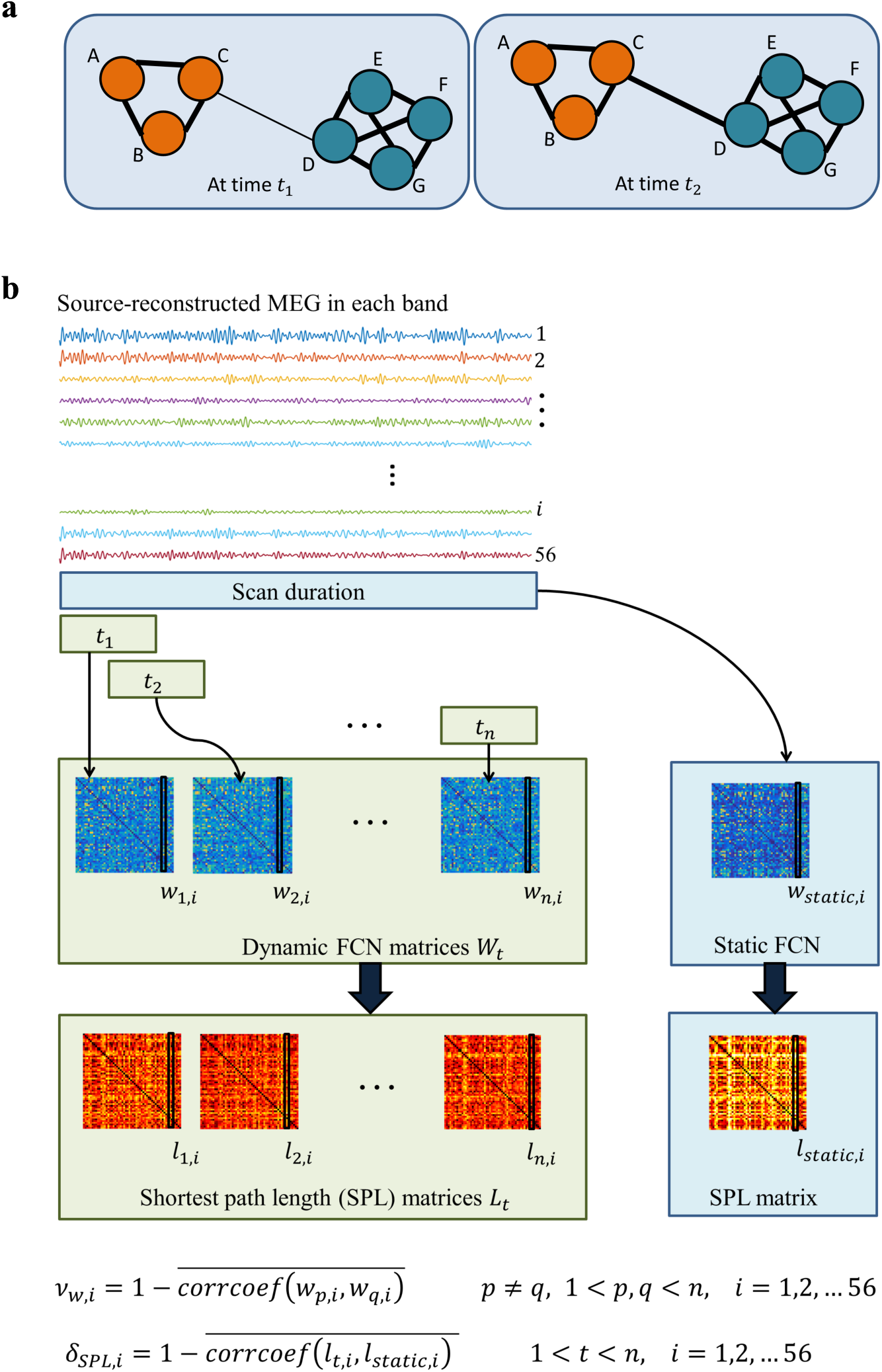
Feature extraction: (a) **Toy example:**Figure shows snapshots of a weighted temporal network(with nodes *A* to *G*) at time instants *t*_1_ and *t*_2_. The thickness of the edges indicate the edge strength. Comparing the graphs, it is seen that the connection strength increases from time *t*_1_ to *t*_2_. Considering only the connections of each node, it is seen that nodes C and D vary between the two instants while others are invariant. However, owing to the topology of the network, the edge (from C to D) also modulates interactions among the other nodes; for example, though A,B are not directly connected by edges to D,E,F,G they can interact with them through C. Hence, any change of edge strength also affects them. In order to capture such variations at the global level which influence temporal dynamics at individual nodes, we propose a measure based on shortest path lengths (see eqn. 3).(b)**Computation of temporal variability:** In each frequency band, the static and dynamic FCNs are estimated from the source-reconstructed MEG. The static connectivity is computed over the entire session while the dynamic connectivity using a sliding window approach using coherence. For computation of dFC, we consider windows of size 2.5s to 15s in steps of 2.5s with 50% overlap between adjacent windows. Using the adjacency matrices of the static and dynamic FCN, we compute the pairwise shortest path lengths between nodes. The temporal variability of *i*^*th*^ region (denoted *δ*_*SPL,i*_) captures the temporal variation of the node with respect to the static network. In addition, *ν*_*w,i*_ Zhang et al. (2016) captures the variations in the functional architecture of the node *i* across time considering its connection strengths to other nodes. However, it may be invariant to global network changes that influence a node’s interactions with others.

In addition to the connection strengths of the node to its neighbours, the role of a node also depends on the role of the nodes it is connected to. Hence, even though a network change does not specifically occur at the node under consideration, it can modify how the node interacts with the rest of the network. To address the drawback of *ν*_*w,i*_, the proposed measure temporal variability (*δ*_*SPL*_) is based on shortest path lengths that take the global network topology into account. Here, the computation of *δ*_*SPL*_ is based on the underlying assumption of connectedness of the graphs being compared. Since we use weighted graphs for determining the shortest paths between nodes, the connectedness of the graph is not an issue. To extend the use of the measure to binary graphs, thresholding the adjacency matrices while retaining the maximum spanning tree may be performed.

### 2.8. Measures of node centrality in static and dynamic FCN

To investigate the relation between the node’s centrality and its temporal variability we quantified the former using the betweennes centrality measure. For a given node *i*, it is computed as the number of shortest paths of the network passing through the node. For each participant, we computed the betweenness centrality of all nodes of the static network and averaged the estimate across sessions, denoted *BC*_*static,i*_ for node *i*. For the dynamic network, we estimated the betweenness centrality of the nodes in each time-window *k*, denoted *BC*_*k,i*_ for node *i*. To characterize the stability of hub structure across time, we averaged the betweenness measure for each node across time. The time-averaged node betweenness for node *i* (denoted ⟨*BC*_*t,i*_⟩) was obtained as:

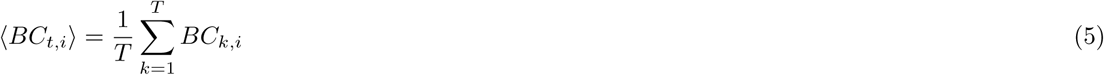

where *T* is the total number of windows. To obtain a single value for time-averaged betweenness for each node per participant, we averaged the values across the multiple sessions and window sizes (2.5, 5, 7.5, 10, 12.5 and 15 sec) for each participant.

### 2.9. Connectome Predictive Modeling

Connectome predictive modeling (CPM) Shen et al. (2017) is a technique developed to predict individual behavior from brain connectivity. The CPM model is built from available training data and makes predictions on novel out-of-sample subjects. To determine whether network variability predicted KABC scores in novel individuals, a leave-one-out cross-validation procedure was adopted. In each fold, features selected from (*n* − 1) participants were used to build the regression model, and the prediction is made on the left out participant. Given the brain connectivity data of a number of participants and target KABC scores, the major steps followed during training are briefly outlined below:

- **Standardized residuals as target scores:** The scores of the training participants on the KABC scale under consideration are regressed on age and gender and the residuals are obtained. For the test data, the residual of the K-ABC scale after regressing out age and gender using the coefficients determined during training. In the following steps, the standardized residuals are used as the target scores for modeling purposes in CPM.
- **Feature extraction:** This step computes relevant features from the brain network of each participant. Features represent a finite, compact representation of the functional connectivity networks (FCN). For example, studies such as Rosenberg et al. (2016); Beaty et al. (2018) have employed the edge strengths in the FCN as features. Here, we used the nodal temporal variability features (*δ*_*SPL*_, eqn. 3) of the participants. Since there are 6 bands and 56 ROI, the feature dimension for a given participant is 6 × 56 = 336.
- **Feature selection:** The correlation between the features and the scores is computed, and those features that are significantly correlated at (*P < α*) are chosen. Since feature selection is performed in every fold, the number of selected features varies from fold to fold. We used the Spearman rank correlation of the features with the scores to perform feature selection with threshold of *α* = 0.01. Further, CPM was performed considering positively and negatively correlated feature subsets seperately.
- **Feature summarization:** The selected features are summarized to obtain a single value per participant in the training data. Here, we used the value of the sum of the features as the summary measure.
- **Model building:** We used simple linear regression model to predict the target scores of the training participants using their summarized features.

The selected features are extracted from the test participant’s connectivity data and the trained model is used to predict the scores on novel participants. The correlation coefficient between the predicted and actual scores was used the performance metric; the higher the correlation the better the predictive capability of the selected features. To quantify the extent to which the features can predict the scores of novel participants, the correlation coefficient between the observed and predicted scores across folds (denoted *ρ*_*CPM*_) was computed. Further, we performed non-parametric permutation tests to evaluate the significance of the predictions. In each iteration of the permutation test, the target scores were shuffled, and the above procedure is followed to compute 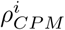, where *i* indexes the *i*^*th*^ iteration. We performed 500 iterations of permutation testing to evaluate the significance. The *P* − value was calculated as the fraction of times 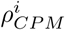, *i* = 1, 2, …, 500 exceeded *ρ*_*CPM*_.

#### 2.9.1. Classification

In addition to CPM which evaluates the relevance of the features for predicting scores (regression), we also considered the problem of correctly predicting the broad *class* a participant belongs to. Specifically, we considered the binary classification problem — *Low* vs *High* for the KABC scales. Given residual scores on any scale (after regressing out age and gender), *Low* and *High* classes were defined for the participants by thresholding the scores; participants below and above the ⅓^*rd*^ and ⅔^*rd*^ quantiles respectively were considered as *Low* and *High*. Leave-one-out cross-validation (LOOCV) was used to evaluate the classification performance, where one participant is tested while pooling all other participants for training. The number of participants considered for classification include (*n* = *n*_*High*_ + *n*_*Low*_), which varies as *n* = 49 for sequential scale and *n* = 51 for simultaneous scale. In each fold, the features from the training data were used to train the classifier and predict the label of the test participant. This was repeated by keeping each participant out for testing. For each considered feature type, Feature selection was performed using the same procedure as in the case of CPM. The selected features^1^ were used to train the classifier. In our experiments, we considered the following linear classifiers:

- **Linear Discriminant Analysis (LDA):** The LDA is a linear classifier which projects data onto a lower-dimensional space, where the decision boundary is determined. For *M* classes, the projection is performed to a *M* − 1 dimensional space, where the direction of projection is chosen to maximize the between-class scatter while minimizing the within-class scatter.
- **Support Vector Machine (SVM):** The SVM is a binary classifier which learns the maximum margin hyperplane in the feature space. It can also be used to learn non-linear decision boundaries by employing the kernel trick. In this case, the features are transformed into a higher-dimensional space and the linear decision boundary is learned. Projecting the decision hyperplane back to the original feature space results in a non-linear decision boundary. We used the SVM classifier with linear kernel and fixed cost parameter *C* = 100.

The overall classification performance of the features was obtained as the average accuracy across folds. The classification performance indicates the proportion of participants classified correctly. Further, the class-wise accuracy indicates the proportion of correctly classified participants from each class.

#### 2.9.2. Empirical chance level

The chance level for a two class classification problem is 50% for a large sample size, but it may be considerably higher for a small data set. Therefore, we estimated the equivalent chance level by taking the sample size into account; assuming that classification errors follow a binomial cumulative distribution Combrisson and Jerbi (2015). The empirical chance level is calculated as the statistically significant decoding accuracy (at *p < α*) for classification accuracy:

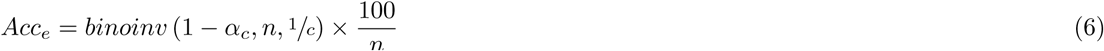

where *n* is the number of samples, *c* is the number of classes, *α*_*c*_ is the threshold for statistical significance.

## 3. Results

In the following, we investigate spatial distribution of temporal variability and age-related differences in dynamic FC of children, and the relation between the node’s variability in the dynamic network and its centrality. To test the role of dynamic FC in cognition, we evaluated connectome predictive modeling Shen et al. (2017) with the KABC scales using nodal temporal variability measures as features, and presented classification performance of the features - i.e., the ability to categorize children as being *Low* or *High* on their cognitive test scores. Finally, we validate our results with alternative measures of nodal temporal variability Zhang et al. (2016).

### 3.1. Temporal variability of brain regions

For each node in the dynamic FCN, we formulated the temporal variability measure denoted *δ*_*SPL,i*_ for node *i* (equation 3) as the dissimilarity of the network architecture within each window with respect to the static network. This captures the network deviations with respect to its manifestation over longer scales (tens of seconds as opposed to few seconds in each window). Hence, higher temporal variability of a node implies that it visits multiple diverse network configurations across time, that are not evident in the static network. In contrast, lower temporal variability suggests that the node maintains close to the intrinsic functional architecture (corresponding to the static network) across multiple time-scales. Also, *δ*_*SPL,i*_ takes into account the fact that even though a network change does not specifically occur at the node *i*, it can modify how the node interacts with the rest of the network (see Figure 1).

#### 3.1.1. Spatial distribution

Figure 2a shows the spatial distribution of temporal variability as measured by *δ*_*SPL,i*_, averaged across all frequency bands and participants. The regions are ranked by their average temporal variability in Figure 2c. The association areas in the posterior parietal and the occipital regions exhibited the highest variability while the temporal and the frontal regions showed much less variability. Figure 2b shows the covariation of *δ*_*SPL,i*_ between nodes, averaged across bands. The matrix entries were computed as the pairwise Spearman rank correlation of the nodes’ temporal variability. We found two distinct clusters of covarying nodes, comprising the nodes in the posterior parietal, occipital regions in one cluster and the frontal, temporal and sub-cortical regions in the other.

**Figure 2:**
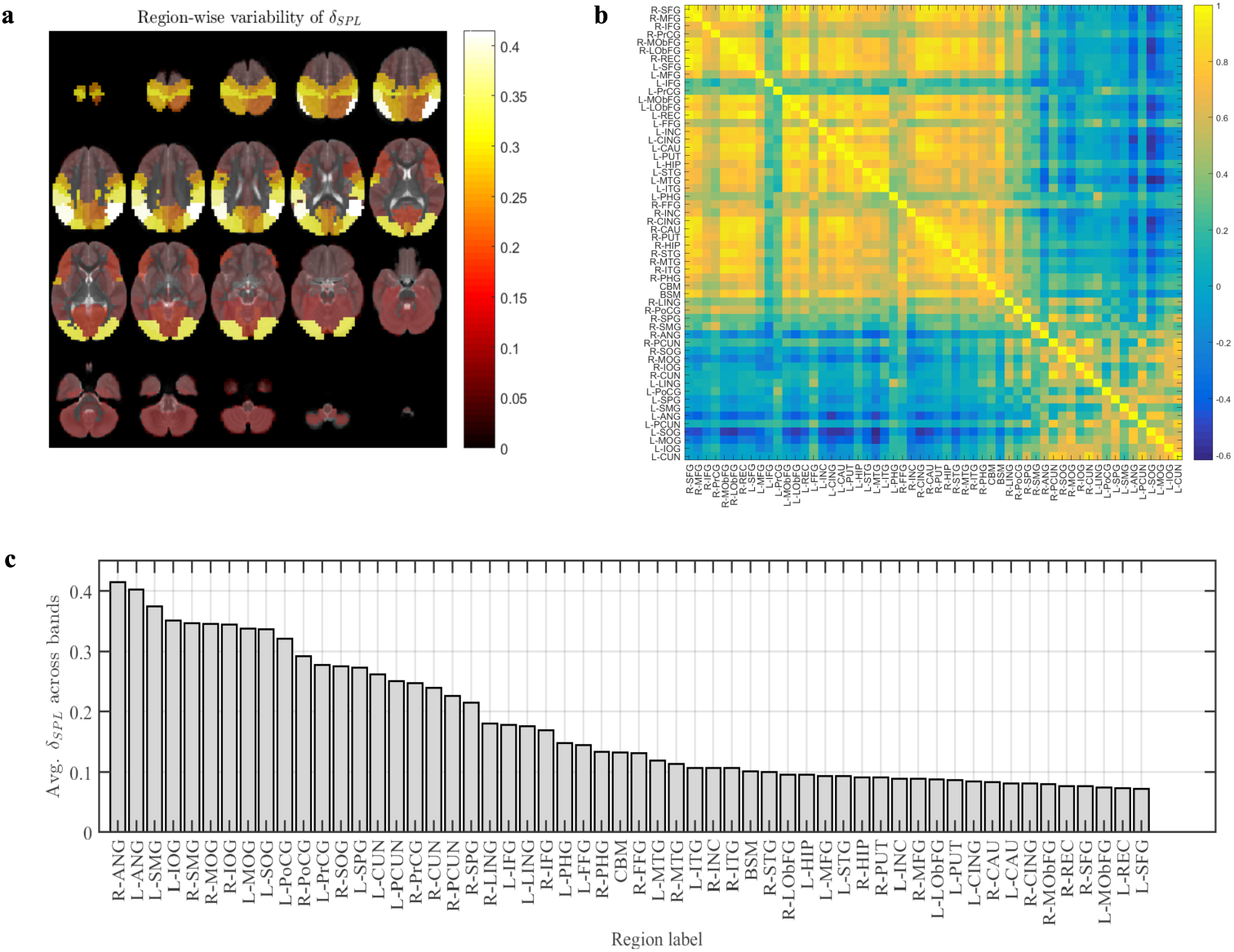
Temporal variability wrt static network: (a) Region-wise variation of the temporal variability measure, *δ*_SPL,*i*_ averaged across all bands (b) Co-variation of temporal variability between node pairs: Figure shows the pairwise Spearman correlation coefficient between the average nodal temporal variability (c)The regions of the brain ordered by their average temporal variability measure in descending order. The full labels of the abbreviations are listed in Supplemental Table S1

#### 3.1.2. Age and gender effects

The overall temporal variability, 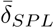, obtained by averaging *δ*_*SPL,i*_ across all nodes and frequency bands, was found to be positively correlated with participant’s age (Spearman *ρ* = 0.3844, *P <* 0.001, *n* = 74), but no significant differences were observed between boys and girls (*t*(74)= 0.753, *P* = 0.454, two-sided). Of note, we observed similar findings in individual frequency bands (see Supplemental Figure S1 and Supplemental Table S2).

### 3.2. Hub structure and temporal variability

To investigate the hypothesis that temporal variability of nodes is associated with their centrality, the latter was estimated using the time-averaged betweenness centrality of the nodes of the dynamic FCN (see equation 5). Figure 3a and 3b show the nodes ranked from the highest to lowest group-averaged betweenness of static (*BC*_*static*_) and dynamic FCN (⟨*BC*_*t*_⟩), respectively. Here the hub nodes (centrality ≥ 1 standard deviation above mean) are found to be similar for the case of both the static and time-averaged betweenness measures. The time-averaged betweenness, ⟨*BC*_*t*_⟩, exhibits nearly perfect linear correlation (Spearman *ρ* = 0.99, *P <* 10^−7^, *n* = 56) with the node-betweenness of the static network, *BC*_*static*_. This suggests that the hub structure in the dynamic network exhibits a stable pattern across time, which is similar to that of the static FCN.

**Figure 3:**
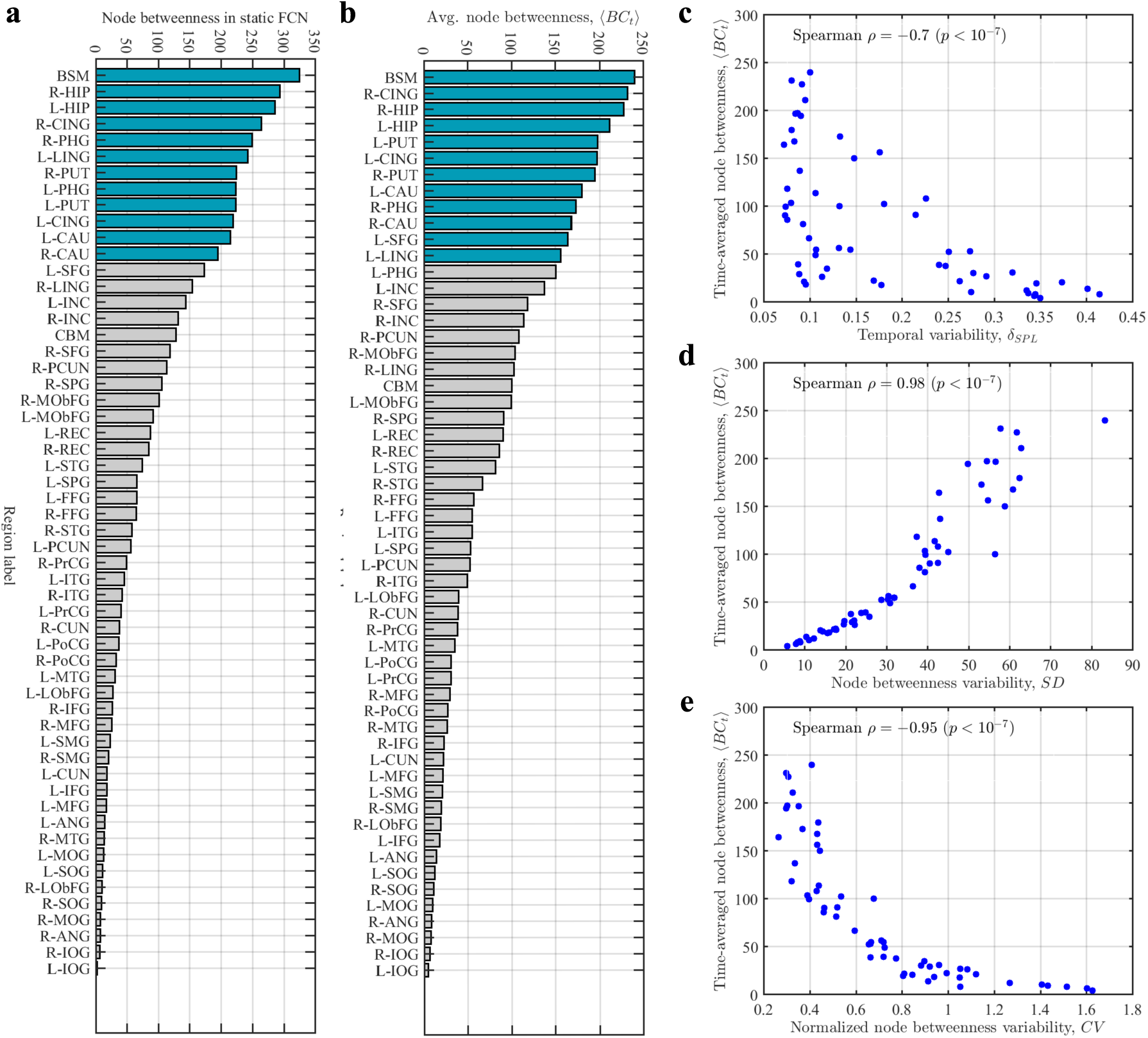
Relation to time-averaged betweenness centrality of dynamic network, ⟨*BC*_*t*_⟩: In the following plots, the group averages are performed across bands and participants. (a) The regions of the brain ordered by their mean *BC*_*static*_ in descending order. The bars in dark cyan are the identified hubs, they represent the nodes scoring ≥ 1 standard deviation above the mean on the centrality measure (b) Same as (a) with ⟨*BC*_*t*_ ⟩ (c) Relation to temporal variability, *δ*_SPL_: The group averaged centrality (⟨*BC*_*t*_⟩) exhibits negative correlation (Spearman *ρ* = − 0.7, *p <* 10^−7^, *n* = 56) with the temporal variability of nodes (d) & (e) relation to variation of node centrality across time (*SD*) and its normalized version *CV* respectively (For a given node *i, SD*_*i*_ and *CV*_*i*_ are computed as the standard deviation and coefficient of variation of time-resolved betweenness *BC*_*t,i*_ across time). The ⟨*BC*_*t*_⟩ measure exhibits positive and negative correlations with *SD* and *CV* respectively. This suggests that though the hub nodes exhibit high variation, this variation is low when compared to their connectivity.

Comparing the ranking of temporal variability (Figure 2c) and betweenness centrality (Figure 3a,b), it is evident that the regions exhibiting high centrality correspond to those exhibiting low temporal variability, and vice versa. This is verified in Figure 3, and the results suggest that temporal variability of the nodes is negatively correlated to their time-averaged betweenness (Spearman *ρ* = − 0.7, *P <* 10^−7^, *n* = 56). This relationship is also consistent in individual bands (see Supplementary Table S3).

However, this observed negative correlation between the hub-structure and the temporal variability is in contrast to earlier studies such as Allen et al. (2012) and Honey et al. (2007) which reported that connections of nodes which exhibit high connectedness (i.e., hubs) tend to fluctuate the most. We suggest that these differences could arise due to the different approaches towards quantifying temporal variability. For example, Honey et al. (2007) measured temporal variability as the extent to which the hub centrality varies across time. To enable comparison, we quantify variation of hub-centrality as the standard deviation of the node-betweenness across time (denoted *SD*_*i*_ for node *i*). In Figure 3d, it is observed that the group-averaged variation in centrality across time is indeed positively correlated to the node centrality (Spearman *ρ* = 0.98, *P <* 10^−7^, *n* = 56). However, normalizing the variation (i.e.,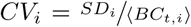) reveals negative correlation with respect to the node centrality (Spearman *ρ* = − 0.95, *P <* 10^−7^, *n* = 56, Figure 3e). Similar observations were also made by Shen et al. (2015), who noted that although hub regions are highly variable in their functional roles at finer timescales, they are not as variable as would be expected by their high centrality.

### 3.3. Connectome Predictive Modeling

To determine whether network variability predicted KABC scores in unseen participants, a leave-one-out cross-validation procedure along with (CPM) Shen et al. (2017) was adopted. In each fold, features selected from (*n* − 1) participants were used to build the regression model, and the prediction was made on the left-out participant. Here, we used the nodal temporal variability features (*δ*_*SPL*_, Eqn. 3) of the participants. Since there are 6 frequency bands and 56 ROIs, the feature dimension for a given participant was 6 × 56 = 336. In each fold, we selected those features which were correlated (Spearman rank correlation, *P <* 0.01) with the target scores. Further, CPM was performed by separately considering positively and negatively correlated feature subsets. We used a simple linear regression model to predict the target scores of the training participants using the selected features. The standardized residuals of the K-ABC scales (after regressing out age and gender) were used as the target scores for modeling purposes in CPM. The selected features were extracted from the test participant’s FCN and the trained model was used to predict the scores on novel participants. The correlation coefficient between the predicted and actual scores was used as the performance metric; the higher the correlation the better the predictive ability of the selected features.

Table 2 shows the results of CPM using temporal variability features, *δ*_*SPL*_ to predict standardized residuals (after regressing out age and gender) of the K-ABC scales. Here, we report the Spearman rank correlation (*ρ*) between the predicted and observed scores across the folds of leave-one-subject-out cross-validation. We assessed the significance of the correlations using *P* − values computed from 500 iterations of non-parametric permutation testing. For the simultaneous and achievement scales,the correlations are not reported since no features were selected for some of the folds and a model could not be built.

**Table 1:** The table shows the details of the participants’ age and K-ABC scores.

| Parameter | Range | mn±s.d |
| --- | --- | --- |
| Age (in months) | 36 – 59 | 48.36 ± 6.99 |
| Sequential scale | 60 – 145 | 98.97 ± 15.56 |
| Simultaneous scale | 53 – 128 | 100.4 ± 14.89 |
| Mental Processing scale | 60 – 132 | 100.1 ± 14.15 |
| Achievement scale | 62 – 156 | 103.4 ± 16.01 |

**Table 2:** Results of Connectome Predictive Modeling with *δ*_*SPL*_ features: The results of CPM were evaluated using leave-one subject-out nested cross-validation (LOOCV), the correlated features (at *p < α*) in each fold were used to predict the score of the test subject using simple linear regression. Table shows the Spearman rank correlations (*ρ* values) between predicted scores and observed scores at two different thresholds of *α* = 0.05, and *α* = 0.01, along with the average no. of selected features across folds, denoted *n*. Here, the *P* −values were computed using non-parametric permutation tests.

| K-ABC scale | Sign of correlation | $n_{Sel}$ | CPM with feature summarization | | CPM without feature summarization | |
| --- | --- | --- | --- | --- | --- | --- |
| | | | Spearman $\rho$ value | $P$ value | Spearman $\rho$ value | $P$ value |
| Sequential scale | Positive | 13.41 | 0.366 | 0.02 | 0.451 | 0.006 |
|  | Negative | 0 | - | - | - | - |
|  | Both | 13.41 | 0.366 | 0.024 | 0.451 | 0.012 |
| Simultaneous scale | Positive | 0 | - | - | - | - |
|  | Negative | 0.04 | - | - | - | - |
|  | Both | 0.04 | - | - | - | - |
| Mental Processing scale | Positive | 5.68 | 0.325 | 0.048 | 0.306 | 0.058 |
|  | Negative | 0.92 | - | - | - | - |
|  | Both | 6.59 | 0.272 | 0.096 | 0.368 | 0.022 |
| Achievement scale | Positive | 0 | - | - | - | - |
|  | Negative | 0.08 | - | - | - | - |
|  | Both | 0.08 | - | - | - | - |

For the sequential scale, the predictions of regression using *δ*_*SPL*_ are significant at *P <* 0.05 (uncorrected −*P* values) for positively correlated features; Figures 4a,b show the relation between the predicted and target scores across folds. The positively correlated features exhibit greater predictive power and larger in number than the negatively correlated ones. The features positively correlated (*P <* 0.01) with the standardized residuals are listed in Table 3.We observed that all the correlated features are from high frequency bands (beta-2 and gamma) and from the left hemisphere; they are primarily from prefrontal cortex (inferior, middle and superior frontal gyri, gyrus rectus, lateral orbitofrontal gyri), temporal areas (inferior, middle, superior temporal gyri), and deeper brain regions (hippocampus, caudate, putamen, insular cortex).

**Table 3:** The table shows the temporal variability features (computed using *δ*_*SPL*_) correlated to the sequential scale at *p <* 0.01.

| Region | Band | Spearman $\rho$ |
| --- | --- | --- |
| L superior frontal gyrus | Beta-2 | 0.3 |
| L middle frontal gyrus | Beta-2 | 0.331 |
| L lateral orbitofrontal gyrus | Gamma | 0.312 |
| L gyrus rectus | Beta-2 | 0.305 |
| L superior temporal gyrus | Gamma, Beta-2 | 0.495, 0.388 |
| L middle temporal gyrus | Gamma | 0.401 |
| L inferior temporal gyrus | Gamma, Beta-2 | 0.481, 0.345 |
| L insular cortex | Gamma, Beta-2 | 0.343, 0.327 |
| L caudate | Beta-2 | 0.299 |
| L putamen | Gamma, Beta-2 | 0.323, 0.318 |
| L hippocampus | Gamma | 0.416 |

**Figure 4:**
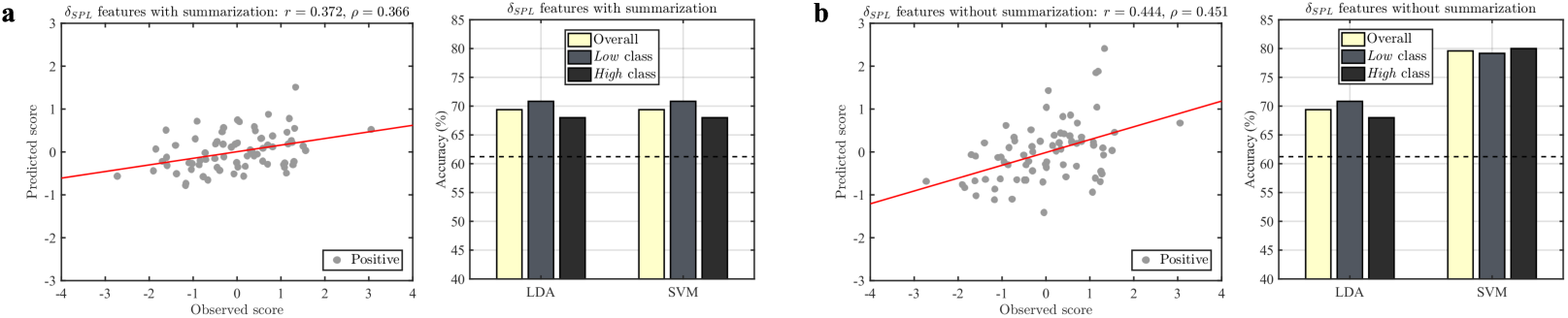
Out-of-sample testing for sequential scale: The results of CPM and binary classification were evaluated using leave-one subject-out nested cross-validation (LOOCV). In CPM, the correlated features (at *p <* 0.01) in each fold were used to predict the score of the test subject using simple linear regression. In addition, the classifier models were built on the training data using correlated features and used to predict the label of the test participant as being *Low* or *High*. The top panel shows results of CPM and classification for temporal variability features (*δ*_*SPL*_) (a) when selected features in each fold are summarized (b) when all selected features were used for model building. The scatter plots show the correlation between the observed and predicted scores on the sequential scale across the folds of LOOCV (both Pearson’s *r* and Spearman’s *ρ* are reported along with the least-squares fit line (red)). The bar plots show the overall and class-wise classification accuracy using two linear classifiers namely, linear discriminant analysis (LDA) and support vector machine (SVM). In the bar plots, the dashed line represents the empirical chance level (see eqn. 6).

For the mental processing scale, the number of positively correlated features also dominated the negatively correlated ones. For the positively correlated features, the relation between the predicted and target scores across folds is shown in Figures 5a,b; the features could weakly predict the scores though the predictions did not attain statistical significance. On the other hand, using both positive and negatively correlated features yielded the best model; relation between the predicted and target scores across folds is shown in Figures 5c,d. We noted that the predictions obtained using multiple selected features in the model (as opposed to summarizing them) yield relatively better predictions. This may be due to complementary information offered by the features, in which case, their sum does not faithfully reflect all the information offered by the features. Table 4 shows the features correlated to the target scores at *P <* 0.01. The positively correlated features are from left temporal areas (inferior, middle, superior temporal gyri), and deeper brain regions (hippocampus, caudate, putamen, insular cortex), while on the other hand, the left parahippocampal gyrus exhibits negative correlation with the scores.

**Table 4:** The table shows the temporal variability features (computed using *δ*_*SPL*_) correlated to the mental processing scale at *p <* 0.01.

| Region | Band | Spearman $\rho$ |
| --- | --- | --- |
| L superior temporal gyrus | Gamma | 0.401 |
| L middle temporal gyrus | Gamma | 0.344 |
| L inferior temporal gyrus | Gamma | 0.412 |
| L parahippocampal gyrus | Beta-1 | -0.322 |
| L insular cortex | Gamma | 0.309 |
| L hippocampus | Gamma | 0.358 |
| cerebellum | Beta-2 | 0.302 |

**Figure 5:**
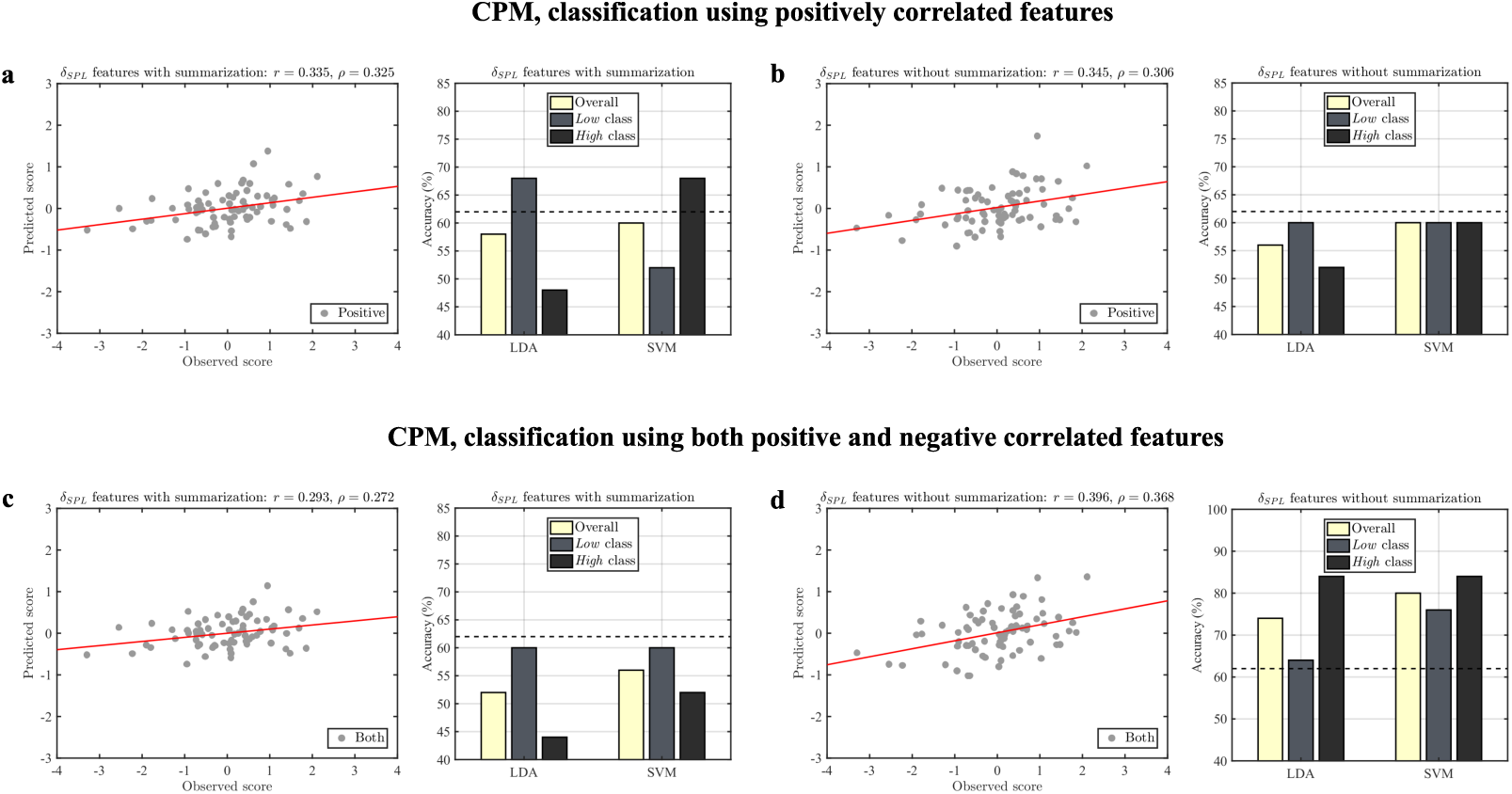
Out-of-sample testing for mental processing scale: The results of CPM and binary classification were evaluated using leave-one subject-out nested cross-validation (LOOCV). In CPM, the correlated features (at *p <* 0.01) in each fold were used to predict the score of the test subject using simple linear regression. In addition, the classifier models were built on the training data using correlated features and used to predict the label of the test participant as being *Low* or *High*. The top panel shows results of CPM and classification for positively correlated temporal variability features (*δ*_*SPL*_) (a) when selected features in each fold are summarized (b) when all selected features were used for model building. The scatter plots show the correlation between the observed and predicted scores on the mental processing scale across the folds of LOOCV (both Pearson’s *r* and Spearman’s *ρ* are reported along with the least-squares fit line (red)). The bar plots show the overall and class-wise classification accuracy using two linear classifiers namely, linear discriminant analysis (LDA) and support vector machine (SVM). In the bar plots, the dashed line represents the empirical chance level (see eqn. 6). Bottom panel, (c) and (d): same as (a) and (b) for for both positively and negatively correlated temporal variability features.

### 3.4. Classification using temporal variability measures

For the sequential and mental processing scales, the features found to significantly predict the scores were also evaluated in a machine learning based classification framework to assess their ability to predict the participant categories (*Low vs High*) on an individual basis. Leave-one-out cross-validation was used to evaluate the classification performance of the selected features (feature selection follows same procedure as in CPM) using linear discriminant analysis (LDA) and support vector machine (SVM) classifiers. The classification accuracy indicates the proportion of participants classified correctly. Further, the class-wise accuracy indicates the proportion of correctly classified participants from each class. A classifier model is deemed useful if the classification accuracy substantially exceeds the empirical chance level Combrisson and Jerbi (2015) (see equation 6), i.e. the expected classification accuracy if the class-labels of the test data were randomly assigned.

Figure 4 shows the results of CPM along with classification performance for positively correlated *δ*_SPL,*i*_ features using LDA, and SVM (linear) classifiers for sequential processing scale. In addition to evaluating the results of classification using sum of selected features (with feature summarization, Figure 2a), we also considered selected features as a vector input to the classifier (without feature summarization, Figure 2b). It is seen that the latter case performs better resulting in classification accuracy (69.387% with LDA, 79.59% with SVM) that substantially exceeds the empirical chance level (61.22%) with balanced class-wise accuracy. Together with the results of CPM, this suggests that the selected features are robust predictors of scores, as well as class labels, on sequential scale in novel or unseen participants.

Figure 5 shows the results of CPM along with classification performance for the mental processing scale for (a,b) positively correlated *δ*_SPL,*i*_ features (c,d) both positively and negatively correlated *δ*_SPL,*i*_ features. It is seen that the only the latter features supplied as vector input to the classifier could significantly predict the class label, yielding classification accuracy of 74% and 80% with LDA and SVM classifiers respectively and exceeds the empirical chance level (62%). This suggests that both positive and negatively correlated network dynamics are important to distinguish low and high-scoring children mental processing scale. Moreover, the inability of the summarized features to classify the participants suggests that the constituent features comprise complementary information.

### 3.5. Comparison to other measures of temporal variability

To validate our results, we also repeated our experiments with the measure of temporal variability proposed in Zhang et al. (2016), computed as *ν*_*w,i*_ for node *i* (see equation 4). The overall temporal variability measured using 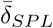 was found to be positively correlated to that measured using 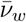 (Spearman *ρ* = 0.794, *P <* 10^−^7). Also, the group-averaged correlation between *δ*_*SPL,i*_ and *ν*_*w,i*_ was statistically significant for all frequency bands even after Bonferroni correction (see Supplementary Table S3), suggesting that both measures show similar patterns of variation across the regions of interest. The regionwise distribution of *ν*_*w,i*_ is shown in Supplemental Figure S2.

The results of connectome predictive modeling using *ν*_*w*_ features for KABC scales (standardized residuals) yielded similar interpretation for sequential scale as *δ*_*SPL*_ features (See Supplemental Tables 2, S5). The results of classification are shown in Supplemental Figure S3. For the mental processing scale, the results of CPM did not attain statistical significance.

## Discussion

In this study, our principal hypothesis was that dynamic functional connectivity network (FCN) would be linked to development and cognitive abilities of preschool children; we focused on preschoolers because it is a critical phase in our development associated with dramatic changes in the brain. We examined dynamic functional brain networks obtained from source reconstructed MEG recordings of 3-4 year old children. We found that the parietal-occipital regions manifested high variability. The overall temporal variability of the functional brain networks increased with age, even within the narrow age-range we considered. At the nodal level, the temporal variability of the brain regions exhibited an inverse correlation with the time-averaged betweenness centrality measure. Importantly, by using connectome predictive modeling and machine learning based classification frameworks revealed that the temporal variability of the brain regions could reliably predict the performance of children’s cognitive abilities, as measured by the K-ABC standardized assessment tests. Finally higher performance on the sequential processing scale was associated with relatively higher variability of regions in the left frontal-temporal areas, while higher performance on the mental processing scale was associated with higher variability of left temporal regions and lower variability of the left parahippocampal gyrus. Taken together, these findings provide critical evidence supporting our hypothesis and further demonstrate that the cognitive abilities can be reliably predicted from a child’s unique dynamical functional connectivity profile.

### Temporal variability of brain regions

We recorded MEG from children while they watched Japanese animation videos and studied the reconfiguration of their functional brain networks across time. Data acquisition during movie watching improves compliance in young children, thus reducing movement artifacts in the recordings. Also, it allows the recording of neural activity during the viewing of naturalistic stimuli Cantlon and Li (2013) and enhances brain-behavior correlations Vanderwal et al. (2018) and test-retest reliability Sonkusare et al. (2019).

In our analyses, we formulated the temporal variability measure (equation 3) of each node in the dynamic FCN to capture the deviations of the node’s role at each time from the static network. This measure is motivated by earlier studies that report measurable differences between static FCN and snapshots of the dynamic FCNAllen et al. (2012); Marusak et al. (2017, 2018). Moreover, any network level change can potentially influence how a node interacts with the rest of the network. For this reason, the proposed measure of temporal variability (*δ*_*SPL*_), adapted from Zhang et al. (2016), is based on the shortest path lengths that take the global network topology into account. It captures variations in the node’s role across time with respect to that in the static network.

Also, the temporal variability was computed using a sliding window with short windows of duration 2.5 to 15 seconds. Compared to fMRI, neuroimaging techniques such as MEG and EEG can probe the fluctuations in dynamic connectivity at faster and behaviorally relevant time scales De Pasquale et al. (2010); de Pasquale et al. (2012); Dimitriadis et al. (2018) and reflect the transitions between FC microstates Koenig et al. (2005); Dimitriadis et al. (2013). Together with fMRI studies, which consider window duration of usually 30 seconds or more (e.g., Zhang et al. (2016); Sun et al. (2018); Li et al. (2017); Keerativittayayut et al. (2018), our results suggest that the dynamics of cortical FC take place at multiple time-scales. Besides, theoretical models of neuronal dynamics simulated such rich structure in the dynamics of cortical FC based on known anatomical connectivity of macaque neocortex Honey et al. (2007); Ghosh et al. (2008); Deco et al. (2009).

Our findings show that parietal and occipital regions of the child brain comprising the inferior parietal lobule (supramarginal, angular gyri) and bilateral visual cortex (inferior, middle, superior occipital gyri), exhibit the highest temporal variability during movie watching. The sensorimotor areas (pre-central, postcentral gyri), medial visual cortex (cuneus, lingual gyrus), and posterior parietal areas (superior parietal gyrus, precuneus) rank next in their variability. The findings suggest greater variability of the primary sensory and unimodal regions involved in sensory functioning. Of note, our observations differ from those in adults Zhang et al. (2016), where primary sensory areas exhibited the least variability across time in the eyes-closed resting state. To account for differences in the choice of measures, we verified the spatial patterns of temporal variability using the measure in Zhang et al. (2016) (see Eq4 in SI Methods) and observed that this did not influence our findings.

Existing studies show that the temporal dynamics of FC during naturalistic viewing differ from resting-state Li et al. (2019). Unlike the resting-state, the viewer engages with the stimuli continuously while watching movies/cartoons, integrating information across time. In this context, the relatively greater variability of the primary sensory regions likely facilitates the neural representation of continually changing visual inputs at short time-scales. Our observations are consistent with investigations of dynamic FC patterns during movie watching in children and adolescents Li et al. (2019).

### Age-related increase in temporal variability

We observed an age-related increase in temporal variability, suggesting that the presence of larger number of functional states may underlie greater variability in older children which might enable higher cognitive functions. Such interpretation is consistent with increased dynamic fluctuations of FC with age Marusak et al. (2017), and positive associations between the number of expressed FC states. Moreover, McIntosh et al. (2008) found that brain signal variability increases with age and that greater variability is correlated with higher cognitive performance.

Of note, the age-group considered in our study is in the range of 36-59 months as we focused only on preschoolers, but this is a narrow age group as compared to Marusak et al. (2017)(7 - 16 year olds) and McIntosh et al. (2008)(8-15 year and 20-33 year olds). Yet, the association between age and temporal variability even in this narrow age group may be attributed to the continuously changing structural network during the preschool years Casey et al. (2005). Early childhood involves rapid neural network development in the brain associated with cognitive and sensory functioning. We suggest that the age-related increase in temporal variability reflects the structural changes in the brain network, that facilitates the dynamic switching among a repertoire of functional states. This possibility is supported by the findings by Tang et al. (2017) that suggest that (structural) brain networks become optimized with age to support diverse brain states. Nevertheless, further studies are needed to determine the nature of influence between structure and function in very young children.

### Relation to hub structure

We characterized the topology of nodes in the static/dynamic FCN using the betweenness centrality. The hubs of the dynamic FCN (identified using time-averaged betweenness centrality) corresponded to the hubs of the static FCN, with a distributed pattern across the brain (sub-cortical structures, superior frontal, lingual gyri, insular cortex, precuneus). Notably, the hub regions overlap with the structural and functional hubs in late childhood (5 - 10 years) Oldham and Fornito (2018). Our observations suggest that dFC microstates are constrained by the underlying structural hubs, which also result in a stable hub-structure across time. This is consistent with recent evidence from fMRI studies in adults, that suggest that dynamic network transitions through various functional metastates whose hubs overlap with that of the structural network Zhao et al. (2019). Further, from a neurodevelopmental perspective, though the structural hubs of the brain are well in place by around 2 years of age Hagmann et al. (2010), the functional hubs shift from being localized in primary sensory and motor areas perinatally to a more distributed pattern in association areas in later years (review in Oldham and Fornito (2018)), reflecting the development of higher-order cognition during this period Casey et al. (2005).

Importantly, we found that the network variability was negatively correlated to the node centrality. This suggests the existence of a core-periphery structure, whose hubs form a stable core and periphery nodes engage/disengage with the core. Such a hypothesis is compatible with the findings of “dynamic core” which found that the hub nodes participate in a number of networks across time facilitating the integration of information de Pasquale et al. (2012); Cole et al. (2013); Schaefer et al. (2014). On the other hand, the periphery nodes lie mainly in the primary sensory areas that bring in the information from the external environment to be interpreted by the brain. The role of peripheral nodes in exploring the dynamical repertoire is also evident in computational studies such as Gollo et al. (2017); here, the authors simulated large-scale fMRI dynamics over known anatomical connectivity of a participant cohort, and found that the local stimulation of periphery regions exhibited larger changes in functional connectivity patterns (of the region with respect to the rest of the brain) as compared to the hub regions.

### Relation to cognitive abilities

The temporal characteristics of functional brain network patterns are robustly related with the scores on sequential and mental processing scales of preschool children, while no such clear relationships are observed for simultaneous and achievement scales. We observed classification accuracy in the range of 70 − 80%, similar to those reported by other developmental studies that used resting state functional connectivity patterns to predict symptom severity Uddin et al. (2013), diagnostic group membership Greene et al. (2016); Chen et al. (2016), and even age Pruett Jr et al. (2015).

In the K-ABC battery, sequential processing involves solving problems with a serial order or sequence of input; for example, the problems in the sequential processing subset are associated with tasks such as word order, number recall and hand movements Kaufman and Kaufman (1983). These tasks require primarily the encoding of information into a short-term memory followed by subsequent recall, therefore, the score on the sequential processing scale is weighted heavily by short-term memory and attention-distractability Bracken (1985); Das (1984). For achieving a high score on the sequential scale, it is essential to maintain multiple items in working memory, against possible interference, for later retrieval and this maintenance phase requires continuously updating the stimuli representation in memory, a process that could be linked to higher network reconfiguration across time. Indeed, this was observed in our findings, and the regions that were positively correlated with this sequential scale were primarily from the auditory and language processing areas (left and right STG, left IFG, MTG), visual processing areas (left and right ITG), left prefrontal cortex (SFG, MFG, ObFG), insular cortex (INS) and sub-cortical regions (CAU, PUT, HIP). Of note, the higher the temporal variability, the more functional communities the brain region will be connected to across time Zhang et al. (2016), thereby allowing a flexible switching between competing network configurations Friston (2000).

Interestingly, our temporal variability measures did not show any clear association with scores on the simultaneous processing scale that requires a holistic or Gestalt approach for integrating inputs to solve problems. The simultaneous processing needs continuous integration of sensory inputs across time, possibly favoring stable network configurations for cognitive processing. Of note, it is emphasized earlier Luria (1976) that the sequential and simultaneous scales are not hierarchical, i.e. one is not more complex than the other Hickman (2008), but see Bracken (1985). Our results therefore emphasize the fact that different mechanisms of dynamic FC underlie sequential and simultaneous information processing, and this would have subsequent implications on accommodating individual differences in children’s learning styles Ayres and Cooley (1986). Further, lower scores on sequential processing scale but with comparable scores on simultaneous processing scale are sometimes associated with children with fragile X syndrome Kemper et al. (1988), ADHD subtype Jonsdottir et al. (2005).

We did observe relationships between temporal variability of FCN and mental processing scale, which combines both sequential and simultaneous processing scales and is a measure of a child’s overall level of cognitive processing McGill and Spurgin (2016). Interestingly, we observed that for the mental processing scale, both positive and negatively correlated network dynamics are important to distinguish between low and high-scoring children. The high scorers on this scale were associated with higher reconfiguration of left temporal brain regions and lower reconfiguration of the left parahippocampal region.

### Limitations and future scope

While this study represents an important role of temporal networks in the cognitive development of very young children, much remains to be explored, both empirically and methodologically, towards a comprehensive understanding of dynamics of functional network and its role in development Kaiser (2017). For example, future longitudinal studies along with model-driven analysis approaches could help revealing the causal role of network variability. Further, here we focused on the neuronal oscillations belonging to classical frequency bands and assumed that the functional network associated with each band to be separated. However, these oscillations are not necessarily independent, instead fast and slow oscillations do interact with each other, enabling a flexible communication and an efficient information transfer between distant brain regions Bonnefond et al. (2017); recent empirical evidence do provide correlated evidence of cross-frequency coupling as a neural measure of intelligence in adults Gągol et al. (2018); Pahor and Jaušovec (2014), and future research should look at the inter-network coupling between interacting neuronal oscillations in children. Further, our MEG data were recorded when the children were watching cartoon of their own choice. This framework of data collection is being increasingly used in neuroimaging studies with young children Richardson et al. (2018), one limitation is that we cannot discriminate between task-driven and intrinsic contributions to the temporal variability of FCN. For example, the negative correlations between left parahippocampal region and mental processing scale might reflect a combination of both intrinsic changes in network structure and increasing stability of the cartoon-driven response in the selected region. Future studies could aim revealing individual contributions of task-driven and intrinsic connectivity by collecting both functional task data and resting state activity from the same child. Finally, we have demonstrated that variability in brain responses seems to be cognitively beneficial at an early stage of development, but excessive variability has specific clinical consequence Dinstein et al. (2015), so it remains an open question whether there exists an optimal degree of reconfiguration ability of functional network in order to support the cognitive flexibility in young children.

## Conclusion

In summary, we provide novel evidence that the dynamical nature of the individual specific functional network topography is refined during development in preschool children and is further linked to their cognitive abilities. Specifically, we found that the children’s scores on a cognitive battery could be predicted to a significant extent using a connectome predictive modeling framework with proposed measures of nodal temporal variability. Further, by using machine learning based cross-validation analysis, we could classify a children as being *Low* or *High*, based on her cognitive score, with an accuracy ranging between 70 − 80%. These findings demonstrate the relevance of temporal networks in establishing brain-to-cognition relationship at a critical phase of early development. Given the increasing evidence of potential links between functional topography and neurodevelopmental disorders Cui et al. (2020); Siugzdaite et al. (2020), we suggest future research to further explore the reconfiguration ability in preschoolers in targeted behavioural interventions Diamond et al. (2007).

## Acknowledgment

The authors L.C. and G.S. were supported under the Signals and Systems for Life Science (SSLS) scheme of Ministry of Human Resource Development, Government of India. The authors would also like to thank Prof. John Richards, University of South Carolina, for granting us access to the Developmental MRI database.

## 4. Supplemental Information

**Table S1:** Names and abbreviations of 56 cortical ROIs

| Region | Abbreviation |
| --- | --- |
| L superior frontal gyrus | L-SFG |
| R superior frontal gyrus | R-SFG |
| L middle frontal gyrus | L-MFG |
| R middle frontal gyrus | R-MFG |
| L inferior frontal gyrus | L-IFG |
| R inferior frontal gyrus | R-IFG |
| L precentral gyrus | L-PrCG |
| R precentral gyrus | R-PrCG |
| L middle orbitofrontal gyrus | L-MObFG |
| R middle orbitofrontal gyrus | R-MObFG |
| L lateral orbitofrontal gyrus | L-LObFG |
| R lateral orbitofrontal gyrus | R-LObFG |
| L gyrus rectus | L-REC |
| R gyrus rectus | R-REC |
| L postcentral gyrus | L-PoCG |
| R postcentral gyrus | R-PoCG |
| L superior parietal gyrus | L-SPG |
| R superior parietal gyrus | R-SPG |
| L supramarginal gyrus | L-SMG |
| R supramarginal gyrus | R-SMG |
| L angular gyrus | L-ANG |
| R angular gyrus | R-ANG |
| L precuneus | L-PCUN |
| R precuneus | R-PCUN |
| L superior occipital gyrus | L-SOG |
| R superior occipital gyrus | R-SOG |
| L middle occipital gyrus | L-MOG |
| R middle occipital gyrus | R-MOG |
| L inferior occipital gyrus | L-IOG |
| R inferior occipital gyrus | R-IOG |
| L cuneus | L-CUN |
| R cuneus | R-CUN |
| L superior temporal gyrus | L-STG |
| R superior temporal gyrus | R-STG |
| L middle temporal gyrus | L-MTG |
| R middle temporal gyrus | R-MTG |
| L inferior temporal gyrus | L-ITG |
| R inferior temporal gyrus | R-ITG |
| L parahippocampal gyrus | L-PHG |
| R parahippocampal gyrus | R-PHG |
| L lingual gyrus | L-LING |
| R lingual gyrus | R-LING |
| L fusiform gyrus | L-FFG |
| R fusiform gyrus | R-FFG |
| L insular cortex | L-INC |
| R insular cortex | R-INC |
| L cingulate gyrus | L-CING |

| **Region** | **Abbreviation** |
| --- | --- |
| R cingulate gyrus | R-CING |
| L caudate | L-CAU |
| R caudate | R-CAU |
| L putamen | L-PUT |
| R putamen | R-PUT |
| L hippocampus | L-HIP |
| R hippocampus | R-HIP |
| cerebellum | CBM |
| brainstem | BSM |

**Table S2:** The relation between *δ*_*SPL*_ and age, gender of participants

| Band | Correlation<br>with age (in months) |  | t-test for age<br>(3- vs 4-year olds) |  | t-test for gender<br>(Boys vs girls) |  |
| --- | --- | --- | --- | --- | --- | --- |
| | Spearman $\rho$ | p-value | t-statistic | p-value | t-statistic | p-value |
| Delta | 0.3190 | 0.0056 | -1.7480 | 0.0847 | 0.2950 | 0.7688 |
| Theta | 0.4050 | 0.0004 | -2.8040 | 0.0065 | 0.9460 | 0.3472 |
| Alpha | 0.3280 | 0.0043 | -1.5520 | 0.1249 | 1.0100 | 0.3161 |
| Beta-1 | 0.0710 | 0.5470 | -0.1900 | 0.8498 | 0.5790 | 0.5641 |
| Beta-2 | 0.3920 | 0.0006 | -3.9890 | 0.0002 | 0.5550 | 0.5809 |
| Gamma | 0.3070 | 0.0078 | -2.3350 | 0.0223 | 0.0070 | 0.9948 |

**Table S3:**
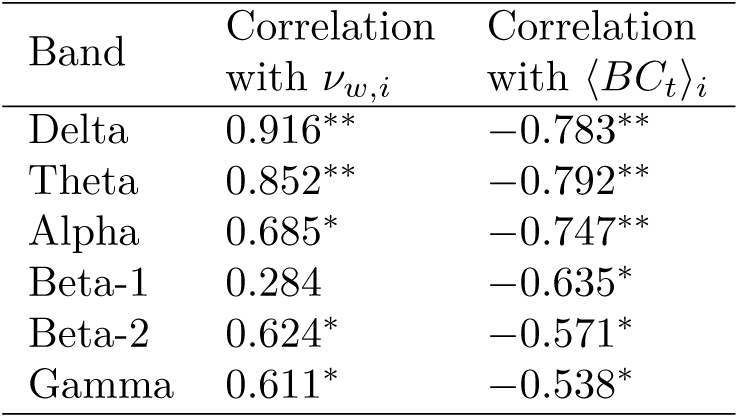
The table shows the Spearman rank correlation between the group-averaged temporal variability measure, *δ*_*SPL,i*_ with *ν*_*w,i*_, ⟨*BC*_*t*_⟩_*i*_ (time-averaged betweenness). Here, ^*\**^ indicates *p <* 0.001,^*\*\**^ indicates *p <* 0.0001

**Table S4:** Results of Connectome Predictive Modeling with *ν*_*w*_ features:

| K-ABC scale | Sign of<br>correlation | $n_{Sel}$ | CPM with<br>feature summarization | | CPM without<br>feature summarization | |
| --- | --- | --- | --- | --- | --- | --- |
| | | | Spearman<br>$\rho$ value | $P$ value | Spearman<br>$\rho$ value | $P$ value |
| Sequential scale | Positive | 21.81 | 0.414 | 0.002 | 0.417 | 0.004 |
|  | Negative | 1.08 | 0.196 | 0.186 | 0.198 | 0.158 |
|  | Both | 22.89 | 0.393 | 0.024 | 0.443 | 0.008 |
| Simultaneous scale | Positive | 0.01 | - | - | - | - |
|  | Negative | 0.07 | - | - | - | - |
|  | Both | 0.08 | - | - | - | - |
| Mental Processing<br>scale | Positive | 7.89 | 0.071 | 0.22 | 0.169 | 0.148 |
|  | Negative | 1.3 | - | - | - | - |
|  | Both | 9.19 | 0.024 | 0.374 | 0.124 | 0.294 |
| Achievement scale | Positive | 0.01 | - | - | - | - |
|  | Negative | 1.8 | - | - | - | - |
|  | Both | 1.81 | - | - | - | - |

**Table S5:** The table shows the temporal variability features (computed using *ν*_*w*_) correlated to the sequential scale at *p <* 0.01.

| Region | Band | Spearman $\rho$ |
| --- | --- | --- |
| L superior frontal gyrus | Gamma, Beta-2 | 0.341, 0.331 |
| L middle frontal gyrus | Beta-2 | 0.384 |
| L middle orbitofrontal gyrus | Beta-2 | 0.428 |
| L lateral orbitofrontal gyrus | Gamma, Beta-2 | 0.37, 0.355 |
| L gyrus rectus | Beta-2, Gamma, Beta-1 | 0.379, 0.334, 0.315 |
| L superior temporal gyrus | Gamma, Beta-2 | 0.365, 0.341 |
| L middle temporal gyrus | Beta-2, Gamma | 0.432, 0.349 |
| L inferior temporal gyrus | Gamma, Beta-2 | 0.327, 0.308 |
| L insular cortex | Gamma, Beta-2 | 0.472, 0.424 |
| L cingulate gyrus | Beta-2 | 0.329 |
| L caudate | Beta-2 | 0.299 |
| L putamen | Beta-2, Gamma | 0.383, 0.355 |
| L hippocampus | Gamma | 0.363 |

**Table S6:** The table shows the temporal variability features (computed using *ν*_*w*_) correlated to the mental processing scale at *p <* 0.01.

| Region | Band | Spearman $\rho$ |
| --- | --- | --- |
| L middle frontal gyrus | Gamma | 0.335 |
| L lateral orbitofrontal gyrus | Gamma | 0.301 |
| L supramarginal gyrus | Beta-2 | 0.306 |
| L superior temporal gyrus | Gamma | 0.32 |
| L middle temporal gyrus | Beta-2 | 0.383 |
| L inferior temporal gyrus | Beta-2 | 0.303 |
| L parahippocampal gyrus | Alpha | -0.328 |
| L insular cortex | Gamma, Beta-2 | 0.35, 0.299 |

**Figure S1:**
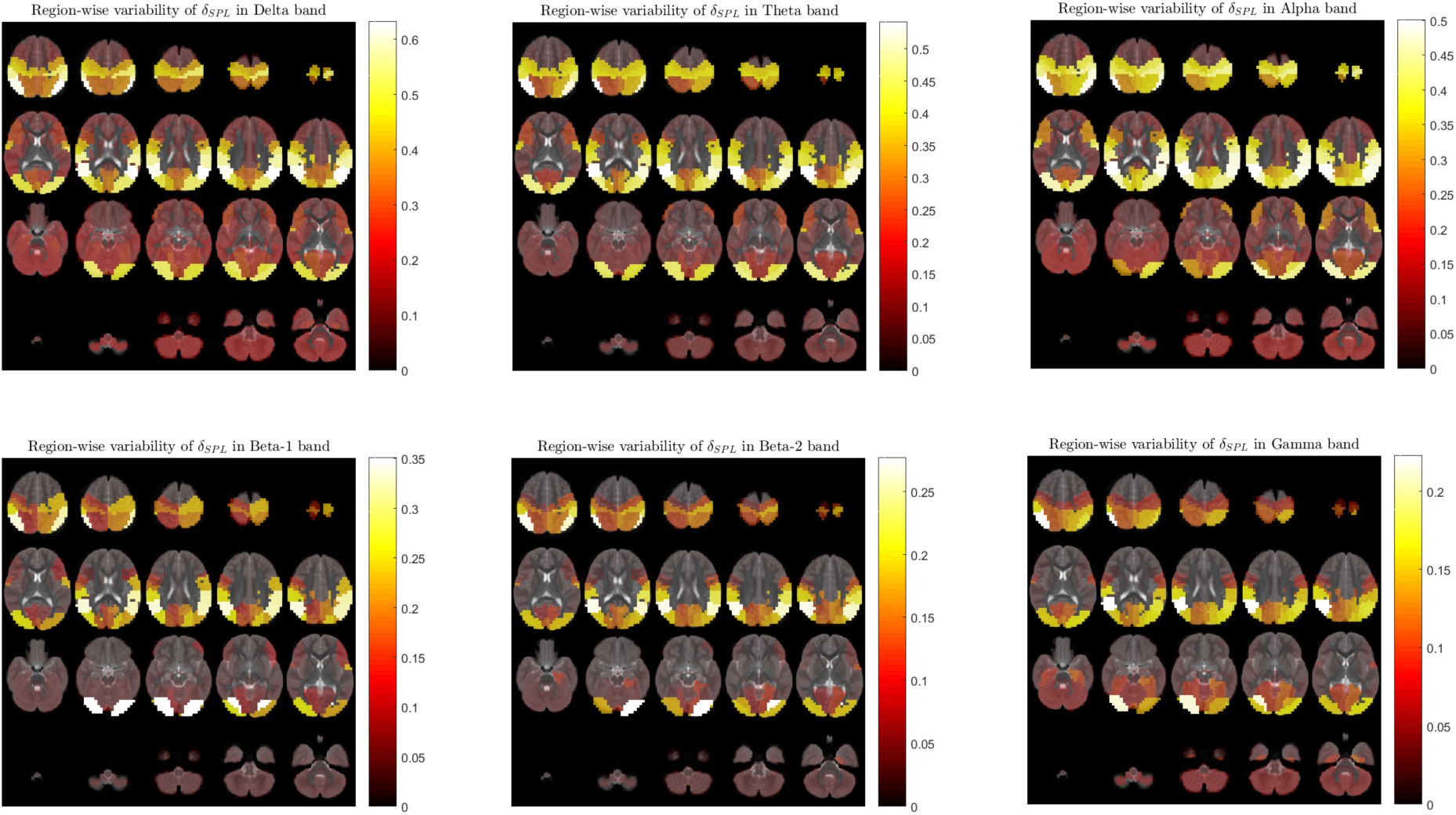
Region-wise variation of the temporal variability measure, *δ*_*SPL,i*_,*i* = 1, 2, …, 56 in individual bands

**Figure S2:**
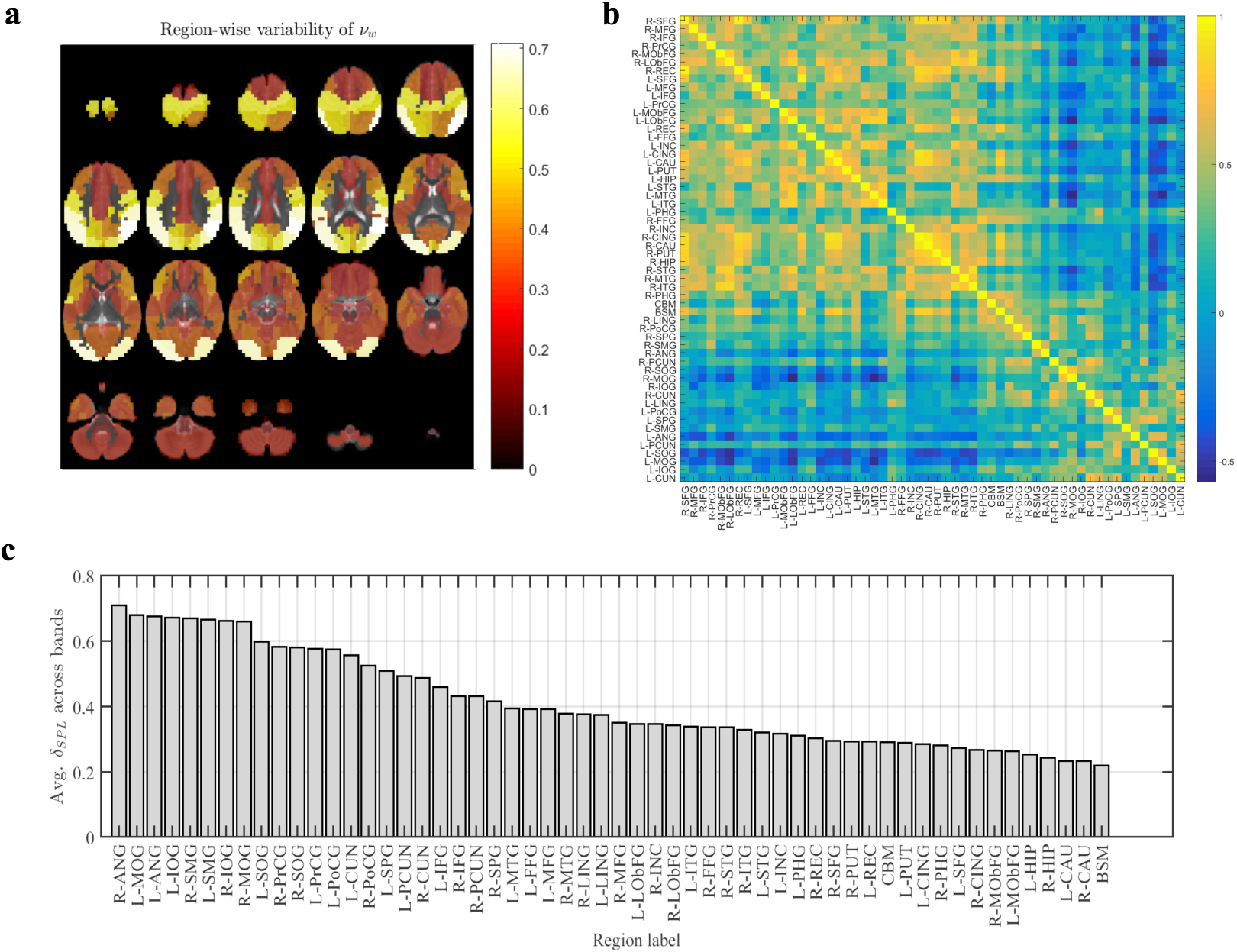
Temporal variability using Zhang et al. (2016): (a) Region-wise variation of the temporal variability measure, *ν*_*w,i*_ averaged across all bands (b) Co-variation of temporal variability between node pairs: Figure shows the pairwise Spearman correlation coefficient between the average nodal temporal variability (c)The regions of the brain ordered by their average temporal variability measure in descending order. The full labels of the abbreviations are listed in Supplemental Table S1

**Figure S3:**
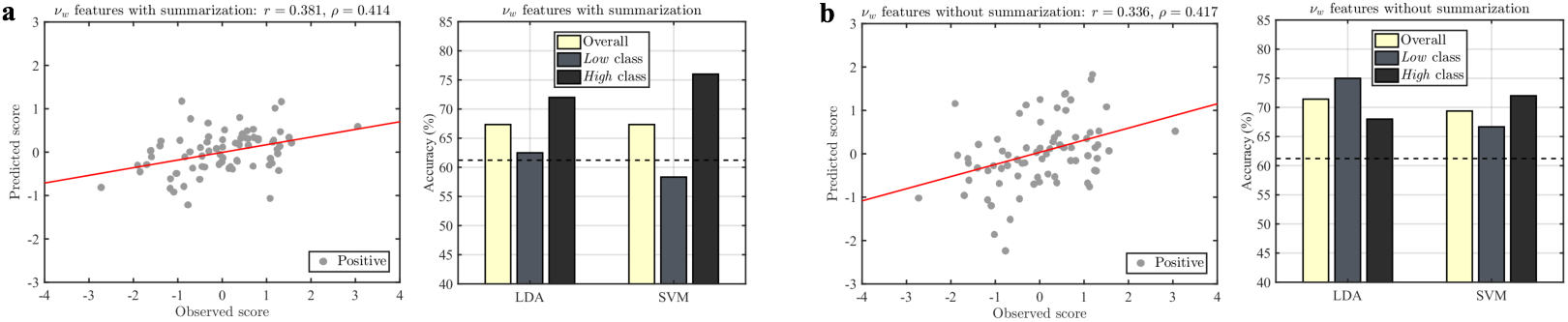
Out-of-sample testing for sequential scale: The results of CPM and binary classification were evaluated using leave-one subject-out nested cross-validation (LOOCV). In CPM, the correlated features (at *p <* 0.01) in each fold were used to predict the score of the test subject using simple linear regression. In addition, the classifier models were built on the training data using correlated features and used to predict the label of the test participant as being *Low* or *High*. The top panel shows results of CPM and classification for temporal variability features (*ν*_*w*_) (a) when selected features in each fold are summarized (b) when all selected features were used for model building. The scatter plots show the correlation between the observed and predicted scores on the sequential scale across the folds of LOOCV (both Pearson’s *r* and Spearman’s *ρ* are reported along with the least-squares fit line (red)). The bar plots show the overall and class-wise classification accuracy using two linear classifiers namely, linear discriminant analysis (LDA) and support vector machine (SVM). In the bar plots, the dashed line represents the empirical chance level (see eqn. 6).

## Footnotes

1 For the binary classification problem, it is to be noted that correlations for feature selection were computed by including the participants with scores in between the ⅓^*rd*^ and ⅔^*rd*^ quantiles along with the participants in the training data from *Low* and *High* classes.

